# Transmission of and susceptibility to seasonal influenza in Switzerland from 2003–2015

**DOI:** 10.1101/252437

**Authors:** Jon Brugger, Christian L. Althaus

## Abstract

Understanding the seasonal patterns of influenza transmission is critical to help plan public health measures for the management and control of epidemics. Mathematical models of infectious disease transmission have been widely used to quantify the transmissibility of and susceptibility to past influenza seasons in many countries. The objective of this study was to obtain a detailed picture of the transmission dynamics of seasonal influenza in Switzerland from 2003-2015. To this end, we developed a compartmental influenza transmission model taking into account social mixing between different age groups and seasonal forcing. We applied a Bayesian approach using Markov chain Monte Carlo (MCMC) methods to fit the model to the reported incidence of influenza-like-illness (ILI) and virological data from Sentinella, the Swiss Sentinel Surveillance Network. The maximal basic reproduction number, *R*_0_, ranged from 1.46 to 1.81 (median). Median estimates of susceptibility to influenza ranged from 29% to 98% for different age groups, and typically decreased with age. We also found a decline in ascertainability of influenza cases with age. Our study illustrates how influenza surveillance data from Switzerland can be integrated into a Bayesian modeling framework in order to assess age-specific transmission of and susceptibility to influenza.

## 1. Introduction

Seasonal influenza has a significant impact on public health. The annual epidemics cause numerous medical consultations and pose a risk for influenza-related complications like viral pneumonia, bacterial superinfection or death in risk groups such as patients with chronic pulmonary or cardiac disease (Taubenberger and Morens, 2008). While the vast majority of influenza-attributed deaths occur in those over 65 years, healthy children under five years have the highest admission rate, particularly infants under six months (Cromer et al., 2014). During pregnancy, influenza infections are a significant and underappreciated public health problem with an increased risk of hospitalization and death (Memoli et al., 2013). In order to better design public health strategies aiming at reducing the burden and morbidity due to influenza, it is indispensable to understand the characteristic transmission patterns of influenza in different populations.

Among the most important parameters that determine influenza transmission in a given population are the basic reproduction number *R*_0_ (i.e., the average number of secondary infections from one infected individual during his or her entire infectious period in a completely susceptible population) and the proportions of specific age groups that are susceptible to the infection. Furthermore, comparing health seeking behavior during different influenza seasons can provide information on the varying severity of the epidemics. These and other critical parameters can be estimated by fitting mathematical models of influenza transmission to epidemiological data (Goeyvaerts et al., 2015; Yuan et al., 2017). A number of studies have systematically analyzed multiple influenza seasons using mathematical models, for example Baguelin et al. (2013) for England and Wales, Lunelli et al. (2013) for Italy, and Goeyvaerts et al. (2015) for Belgium. Only a few studies have used mathematical models to study influenza transmission in Switzerland. Chowell and colleagues analyzed the 1918 influenza pandemic in Geneva (Chowell et al., 2006, 2007; Rios-Doria and Chowell, 2009), and Smieszek et al. (2011) used a spatial individual-based model to study the spread of H3N2 during the 2003/2004 season. To our knowledge, however, there are no mathematical modeling studies that analyze the transmission dynamics of influenza in Switzerland over multiple seasons.

Many countries maintain extensive influenza surveillance systems for tracking the course and extent of the yearly epidemics. In Switzerland, the monitoring is performed by Sentinella, the Swiss Sentinel Surveillance Network, since 1986 (Somaini et al., 1986). This network is a co-project between the Swiss Federal Office of Public Health (SFOPH) and 150–250 general practitioners (GP) who report all cases of influenza-like-illness (ILI) on a voluntary basis. ILI is a symptom complex consisting of typical symptoms of influenza infections, such as malaise, fever, cough, and muscle pain. On the basis of the data collected in this sentinel network, the SFOPH publishes the weekly incidence of ILI-related GP consultations. In addition, some of the patients within the network who suffer from ILI are virologically tested through a nasopharyngeal swab that can be used to determine strain-specific positivity for influenza.

In this study, we conducted a detailed analysis of the transmission dynamics of ten influenza seasons in Switzerland from 2003-2015. We used a compartmental influenza transmission model with seasonal forcing taking into account age-specific social mixing and health-care seeking behavior. We fitted the model to ILI and virological test data from Sentinella in a Bayesian framework using Markov chain Monte Carlo (MCMC) methods. This allowed us to obtain a comparative analysis of the transmissibility of and susceptibility to past influenza seasons in Switzerland.

## 2. Methods

### 2.1. Data

We used data from Sentinella, the Swiss Sentinel Surveillance Network (http://www.sentinella.ch) (Somaini et al., 1986) that were provided by the SFOPH. The data set provides the numbers of ILI-related GP consultations, *z*_*i*_(*n*), per 100,000 inhabitants during week *n*, where *i* = 1,…, 5 indicates five age groups of 0–4, 5–14, 15–29, 30–64, and 65+ year olds. From ISO week 39 to week 16 in the following year, a subset of these patients with ILI were virologically tested for influenza via a nasopharyngeal swab (Hôpitaux Universitaires Genève, accessed 24 Nov, 2017). The swabs were analyzed using viral cell culture until 2005/2006 and using a more sensitive RT-PCR since then. We denoted the total number of virological swab tests during week *n* and for age group *i* as *v*_*i*_(*n*), and the number of tests that were positive for influenza as 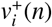. No age-specific virological data was available for the seasons 2007/2008 and 2013/2014, and we excluded these two seasons from our analysis.

### 2.2. Transmission model

We developed a deterministic, population-based model that describes human influenza transmission across different age groups in Switzerland. Assuming an SEIR (susceptible-exposed-infected-recovered) structure and gamma-distributed latent and infectious periods (Keeling and Rohani, 2008), the model can be described by the following set of ordinary differential equations (ODEs):

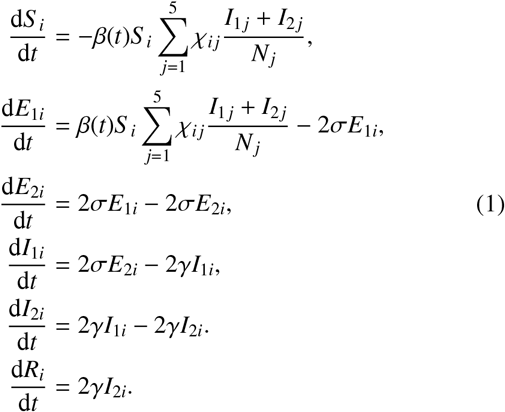

where *i* = 1,…, 5 indicates the five age groups 0–4, 5–14, 15–29, 30–64, and 65+ year olds. We assumed a fixed population size *N* = 100,000, partitioned into the different age groups according to the age distribution in Switzerland (Swiss Federal Statistical Office, accessed 24 Nov, 2017). Individuals are considered susceptible *S* if they have not been infected and have no (cross-)immunity from previous influenza infections or vaccination. After infection, exposed individuals *E* remain latently infected for an average of 1/σ days before they become infectious individuals *I* for an average of 1/*γ* days. After natural clearance, individuals enter the recovered compartment *R*. *β*(*t*) denotes the time-dependent transmission rate (see *Seasonal transmissibility)* and *χ* = (*χ*_*ij*_)_*i,j*_ describes the contact matrix (see *Contact matrix*).

#### 2.2.1. Contact matrix

In absence of population-based survey data (Mossong et al., 2008), the structure of social contacts can be inferred from census and demographic data, such as household size and composition, age structure, rates of school attendance, etc. (Fumanelli et al., 2012). Based on these data, Fumanelli et al. (2012) simulated a population of synthetic individuals in order to derive contact matrices for various member states of the European Union, Norway and Switzerland. We transformed the published matrix of adequate contacts in Switzerland by dividing the matrix by the age structure of the Swiss population (Swiss Federal Statistical Office, accessed 24 Nov, 2017). This resulted in the contact matrix *χ*_*ij*_ that provides the average number of adequate contacts an individual of age group *i* has with individuals of age group *j* (see Supplementary material).

#### 2.2.2. Seasonal transmissibility

Whereas influenza epidemics in tropical and subtropical regions often occur twice a year during the rainy seasons, there is a strong seasonal cycle in temperate regions (Tamerius et al., 2013). The oscillation in transmissibility is most likely caused by changes in temperature and humidity (Shaman et al., 2010, 2011; Lofgren et al., 2007). The sinusoidal curve *β*(*t*) = *β*_0_ + *∊*cos(2*π*(*t* - *ϕ*)/52.14) provides a reasonable approximation to model the seasonal forcing of influenza, where *β*, *∊*, and *ϕ* are auxiliary variables. We used the following parameter transformations to introduce the basic reproduction number 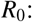 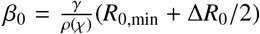 and 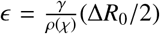, where *R*_0,min_ is the minimum of *R*_0_ and Δ*R*_0_ = *R*_0,max_ − *R*_0,min_. We calculated 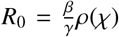 using the next generation matrix (Odo Diekmann, 2013; Heffernan et al., 2005), where *ρ*(*χ*) is the spectral radius of the contact matrix *χ*.

#### 2.2.3. Likelihood function

In order to embed the incidence of ILI and the virological data into a likelihood model, we followed a similar method to the one described by Baguelin et al. (2013). We assumed that for each age group *i* only a proportion *p*_*ai*_ of influenza cases is ascertainable, which means that the following conditions must be fulfilled:

1. the individual is infected with influenza
2. the individual is symptomatic and seeks a GP
3. the GP records the individual as ILI
4. if the GP performs a nasopharyngeal swab, the test result is positive.

While the majority of these ascertainable influenza cases are caused by the national epidemic, there is an additional influx of cases from abroad. We modeled this influx with the constant parameter *ζ*_*c*_. We then described the total incidence of weekly ascertainable influenza cases per 100,000 as follows:

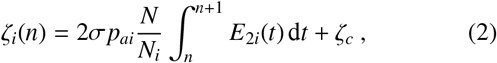

where *n* denotes the corresponding week. We introduced the random variable 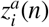 that describes the sampled ascertainable influenza cases according to a truncated negative binomial distribution with dispersion parameter Φ:

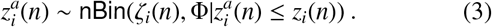

The negative binomial distribution can account for variation in the sampling process, e.g., GP consultations that are not uniformly distributed throughout a week, and additionally allows for over-dispersion of cases due to stochastic processes that are not captured by the deterministic model. We used the following parameterization: if *X* ~ nBin(*μ*, *Ψ*) then E(*X*) = *μ* and Var(*X*) = *μ* + *μ*^2^/*Ψ*. The variable 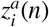 serves as an auxiliary variable, since the ascertained cases cannot be directly derived from the incidence of ILI. Rather, we used 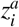 to calculate the proportion 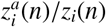, which describes the probability of detecting influenza using a nasopharyngeal swab within the set of ILI-related GP consultations. We can then describe the total number of influenza-positive cases, 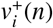, among *v*_i_(*n*) individuals that provided a swab test in age group *i* using a binomial distribution:

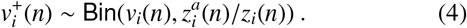

#### 2.2.4. Parameter priors

We used the same prior distributions for the parameters for all ten seasons. The prior distributions for the latency and infectious periods were based on estimates from experimental data. While Cori et al. (2012) described these periods using shifted Weibull distributions, we used gamma distributed durations to accommodate multiple compartments in our transmission model. The prior distributions of *R*_0,min_ and Δ*R*_0_ were informed by findings from Shaman et al. (2010, 2011). We assumed a tight prior distribution for the forcing parameter *ϕ*, which describes the time point when *R*_0_ reaches its maximum, around the first week of the year. This is in agreement with findings from Shaman et al. (2010, 2011) and the time point when absolute humidity is typically lowest in Switzerland (Swiss Federal Office of Meteorology and Climatology MeteoSwiss, accessed 24 Nov, 2017). We set the prior distribution of the probability of ascertainability around 3% for all age groups, which is in agreement with findings from Baguelin et al. (2013). Other parameters, including the susceptibility of different age groups (see *Implementation*) were uniformly chosen within reasonable intervals. Table 1 provides a summary of all free model parameters together with their prior distributions.

**Table 1.** Definitions, prior distributions and posterior estimates of the free model parameters that describe the 2009/2010 influenza epidemic in Switzerland.

| Parameter | Description | Prior | Estimate (median and IQR) |
| --- | --- | --- | --- |
| $1/\sigma$ | Latency period (days) | $\mathcal{N}(1.63, 0.06^2)$ | 1.67 (1.626–1.704) |
| $1/\gamma$ | Infectious period (days) | $\mathcal{N}(0.99, 0.25^2)$ | 1.47 (1.331–1.600) |
| $R_{0,\min}$ | Minimal basic reproduction number | $\mathcal{N}(1, 0.25^2)$ | 1.17 (1.109–1.241) |
| $\Delta R_0$ | $R_{0,\max} - R_{0,\min}$ | $\mathcal{N}(1, 0.25^2)$ | 0.48 (0.417–0.535) |
| $\phi$ | ISO week where $R_0 = R_{0,\max}$ | $\mathcal{N}(1, 1^2)$ | 1.23 (0.572–1.831) |
| $t_0$ | ISO week of simulated onset of influenza season | $\mathcal{U}([25, 52 + 3])$ | 31.89 (28.939–33.889) |
| $\zeta_c$ | Ascertainable influx from abroad (per 100 000) | $\mathcal{U}([0, 5])$ | 0.90 (0.697–1.175) |
| $\Phi$ | Dispersion parameter | $\mathcal{U}([0, 100])$ | 7.38 ( 5.300–10.648) |
| $p_{a1}$ | Ascertainability in age group 0–4 | $\mathcal{N}(0.03, 0.01^2)$ | 0.05 (0.045–0.053) |
| $p_{a2}$ | Ascertainability in age group 5–14 | $\mathcal{N}(0.03, 0.01^2)$ | 0.05 (0.050–0.058) |
| $p_{a3}$ | Ascertainability in age group 15–29 | $\mathcal{N}(0.03, 0.01^2)$ | 0.06 (0.052–0.061) |
| $p_{a4}$ | Ascertainability in age group 30–64 | $\mathcal{N}(0.03, 0.01^2)$ | 0.03 (0.025–0.034) |
| $p_{a5}$ | Ascertainability in age group 65+ | $\mathcal{N}(0.03, 0.01^2)$ | 0.03 (0.020–0.033) |
| $p_{s1}$ | Susceptibility in age group 0–4 | $\mathcal{U}([0, 1])$ | 0.94 (0.882–0.976) |
| $p_{s2}$ | Susceptibility in age group 5–14 | $\mathcal{U}([0, 1])$ | 0.88 (0.817–0.929) |
| $p_{s3}$ | Susceptibility in age group 15–29 | $\mathcal{U}([0, 1])$ | 0.96 (0.933–0.984) |
| $p_{s4}$ | Susceptibility in age group 30–64 | $\mathcal{U}([0, 1])$ | 0.88 (0.790–0.931) |
| $p_{s5}$ | Susceptibility in age group 65+ | $\mathcal{U}([0, 1])$ | 0.30 (0.204–0.408) |

#### 2.2.5. Implementation

We fitted the model to the data for each influenza season individually. At *t*_0_, we initialized the ODEs with one exposed individual partitioned across the five age groups according to their size, i.e., *E*_1*i*_(0) = *N*_*i*_/*N*. We further introduced the parameter *p*_*Si*_ to account for the proportion of susceptible individuals in age group i, i.e., *S*_*i*_(0) = *p*_*si*_*N*_*i*_ − *N*_*i*_/*N* and *R*_*i*_(0) = (1 − *p*_*Si*_)*N*_*i*_. All other compartments were set to zero.

We implemented the MCMC simulations in Stan (Carpenter et al., 2017; Stan Development Team, 2016), a programming language written in C++ using a Hamiltonian Monte Carlo (HMC) procedure with fast convergence. For every season, we sampled two chains of length 1,000 with a burn-in of 500 using UBELIX (http://www.id.unibe.ch/hpc), the HPC cluster at the University of Bern. We visually confirmed convergence using the chain plots together with package `ggmcmc` of the programming language R (R Core Team, 2015). Since Stan does not support sampling of discrete variables, we discretized the likelihood function resulting in the following equation:

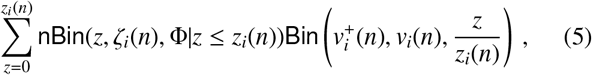

where the overall log-likelihood function is the sum of the logarithms of Eq. 5 over all *i* and *n*. We ignored data points where *v*_*i*_(*n*) = 0 or *z*_*i*_(*n*) = 0. For the truncated negative binomial, we had to calculate the cumulative distribution. We used a normal approximation according to Camp-Paulson to speed up computation in Stan (Bartko, 1966).

## 3. Results

The influenza transmission model is capable to reproduce the seasonal epidemic curves of ascertainable influenza infections for all age groups (Fig. 1). The model fits particularly well to the data from the three oldest age groups (15–29, 30–64, and 65+ year olds), while the numbers of ascertainable influenza infections for the two younger age groups (0–4 and 5–14 year olds) seem to be somewhat underestimated. This trend is particularly apparent for 0–4 year olds where the model underestimates the peak incidence. This systematic trend can be explained by the relatively low number of virological samples for this age group, which leads to a higher uncertainty of the data and a corresponding lower impact on the overall likelihood function during the model fit.

**Fig. 1.**
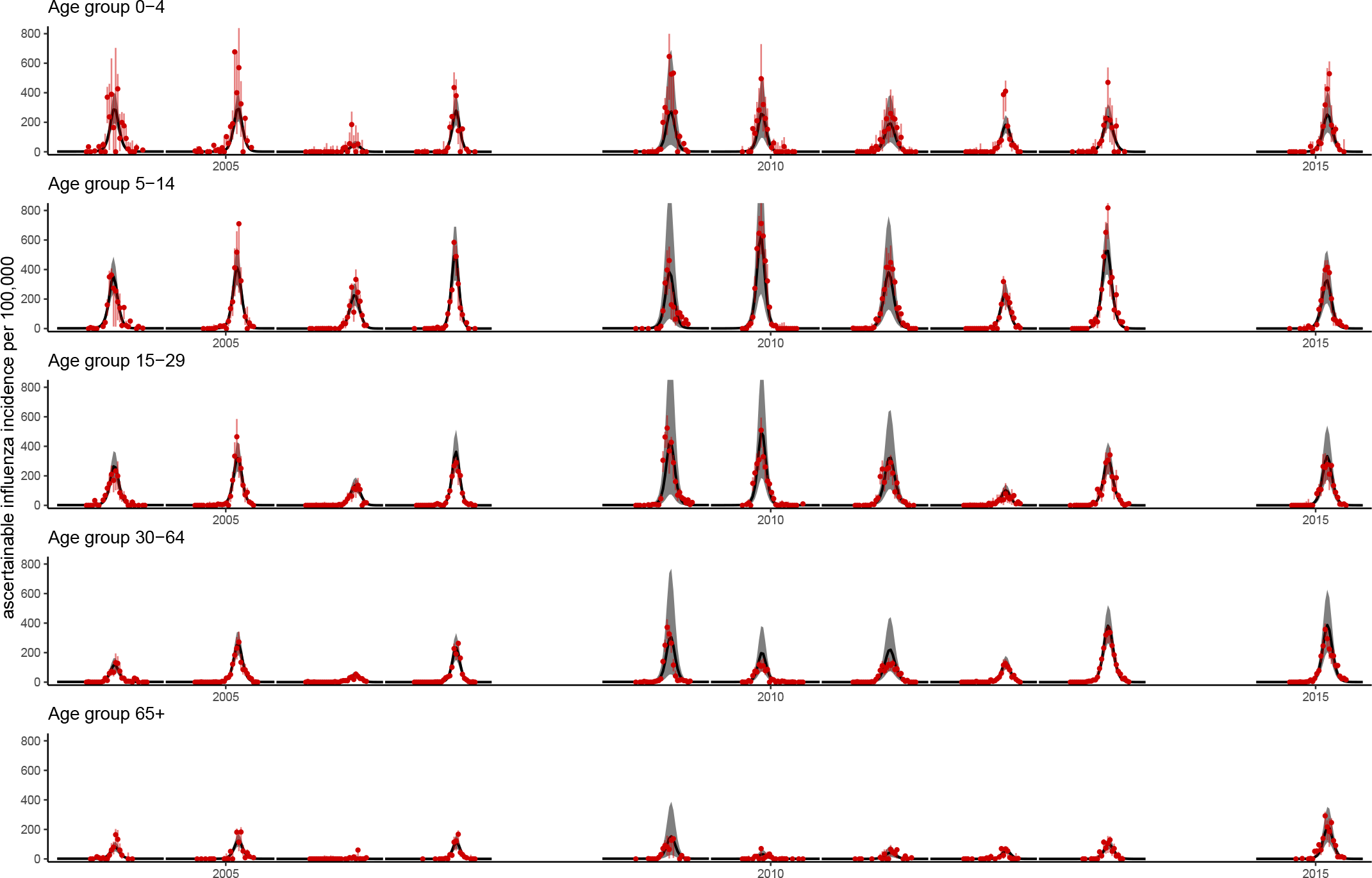
Model fits to data from seasonal influenza epidemics for five age groups in Switzerland from 2003 to 2015. Black lines represent the best-fit model together with 95% credible intervals (gray shaded area). The red dots represent the incidence of ascertainable influenza infections multiplied by the proportion of virological samples that are positive for influenza. The red vertical lines correspond to the 95% confidence intervals of these data according to a binomial distribution. Data are from the Swiss Sentinel Surveillance Network, Sentinella. The influenza seasons 2007/2008 and 2013/2014 were excluded due to the lack of virological data.

The Bayesian modeling framework allowed us to obtain posterior distributions of several parameters that describe seasonal influenza epidemics in Switzerland (Table 1, Fig. 2 and Supplementary Material). While most parameters do not seem to be correlated, we typically found a slight negative correlation between *R*_0,min_ and Δ*R*_0_, between *R*_0,min_ and susceptibility, *p*_*Si*_, and between susceptibility, *p*_*Si*_, and ascertainability, *p*_*ai*_ (see Supplementary material).

**Fig. 2.**
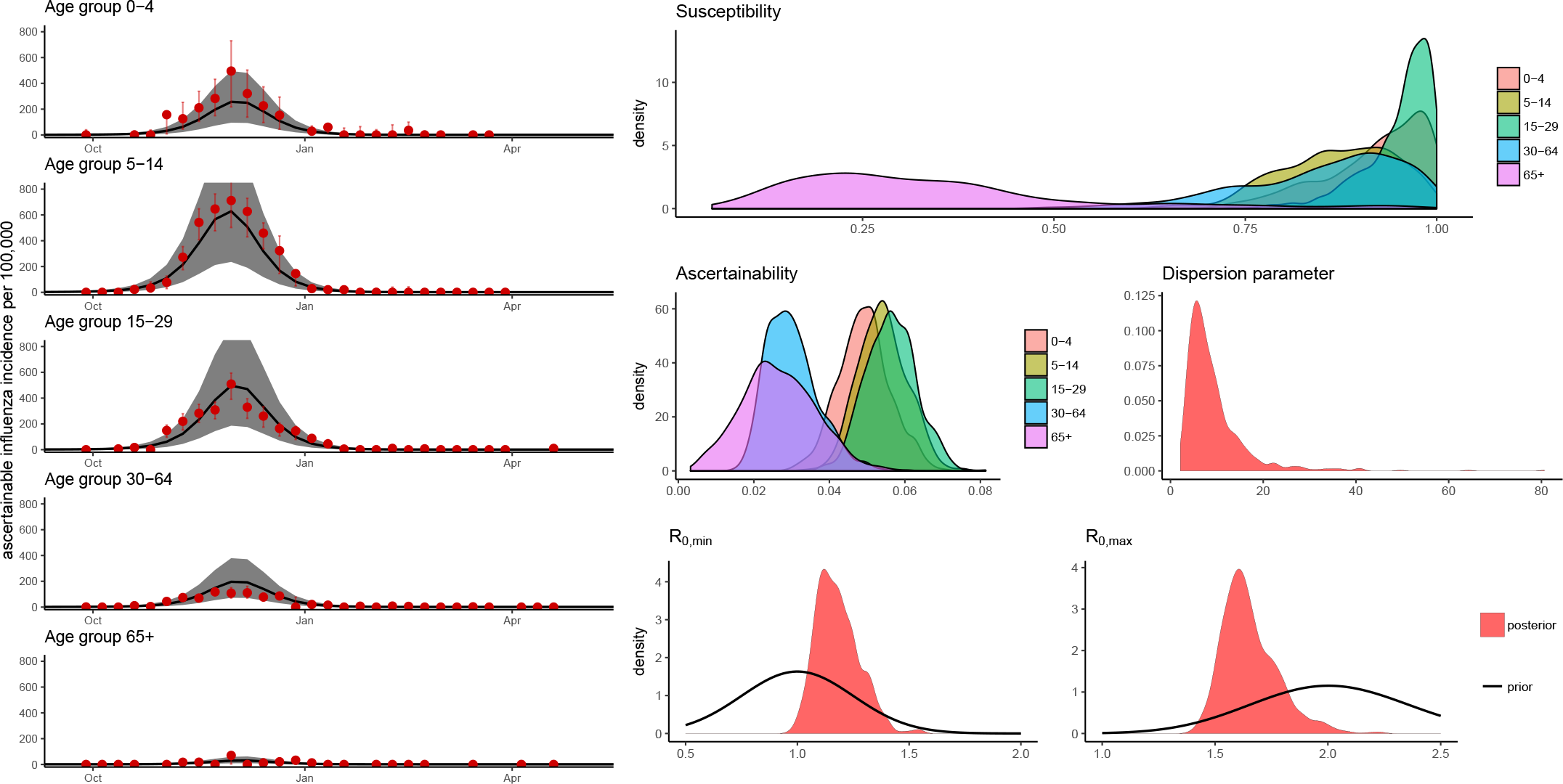
Model fits and parameter inference for the 2009/2010 influenza season in Switzerland. The left panels represent a close-up from Fig. 1. The middle and right panels show posterior distributions of key model parameters. Data are from the Swiss Sentinel Surveillance Network, Sentinella.

Comparing the posterior distributions of some key parameters between all epidemics sheds light on the between-season variability (Fig. 3). The median values of *R*_0,min_ range between 0.91 and 1.50, and the median values of *R*_0,max_ range between 1.46 and 1.81. The pattern of age-specific susceptibility to influenza, *p*_*Si*_, appears to be similar across seasons. Susceptibility is usually highest among 0–4 year olds (seasonal medians between 0.75 and 0.97) and decreases with increasing age. The posterior distributions for susceptibility are typically widest for the two oldest age groups (30–64 and 65+ year olds), which can be explained by the wider age range compared to the younger age groups. The greatest exception from the pattern of decreasing susceptibility with age represents the influenza season 2014/2015. Here, 14–29 year olds show the lowest susceptibility while it is close to 100% for all other age groups. Ascertainability of influenza infections shows a similar pattern to susceptibility and decreases with increasing age, with most median values ranging around 5%. The influenza season 2014/2015 again shows a divergent pattern with the highest ascertainability in the oldest age group. Finally, the median values of the dispersion parameter Φ range from 2.36 in 2008/2009 to 53.18 in 2012/2013. Those seasons with lower values of Φ exhibit higher variability in incidence and are indicative of within-season variability in ascertainability of influenza cases.

**Fig. 3.**
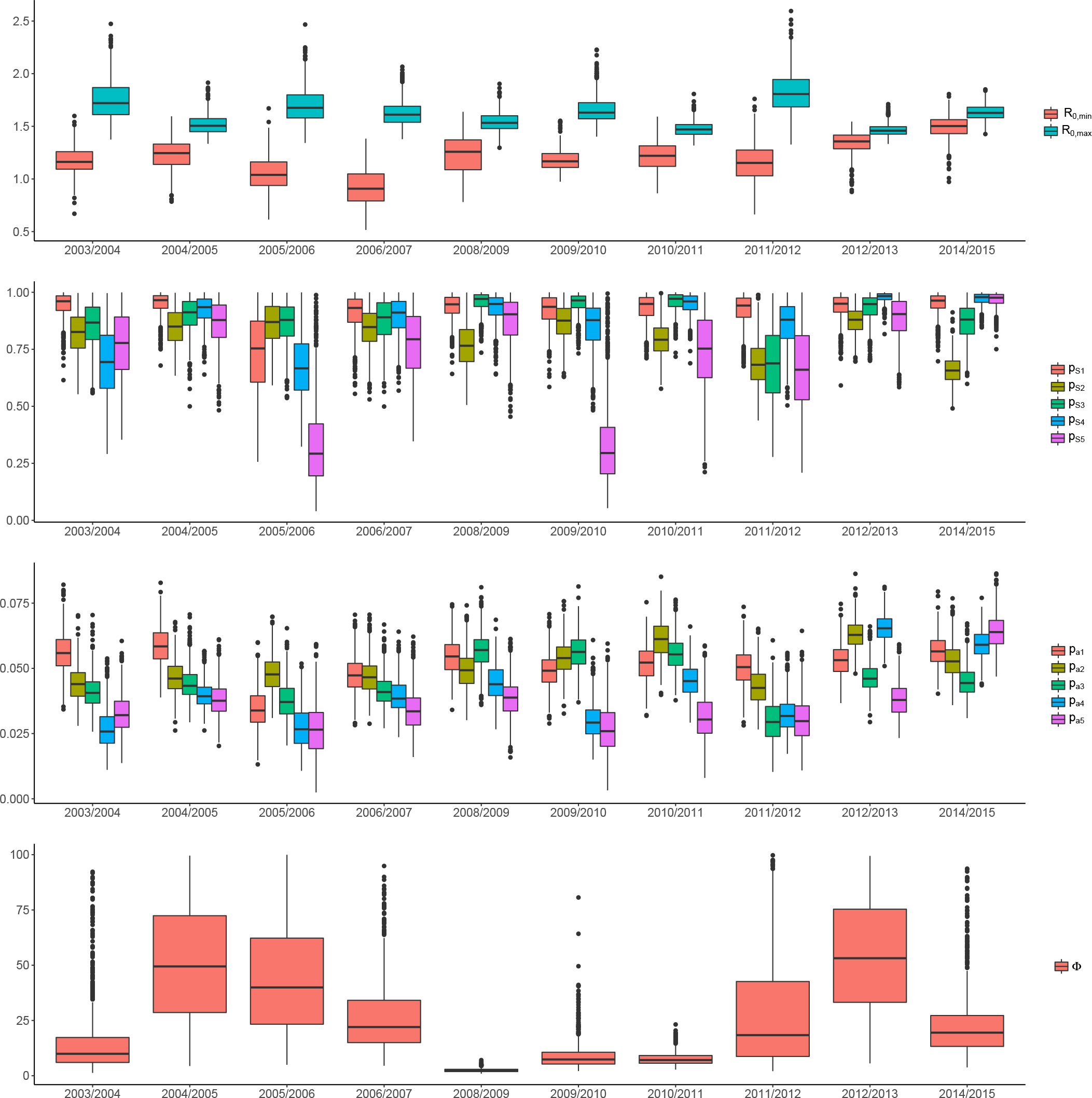
Comparison of key model parameters between influenza epidemics in Switzerland from 2003-2015. Panels show boxplots with interquartile ranges for the minimal (*R*_0,min_) and maximal (*R*_0,min_) basic reproduction number (top), susceptibility (*p*_*Si*_) and ascertainability (*p*_*ai*_) (middle), and the dispersion parameter Φ (bottom).

Since many mathematical modeling studies focus on the H1N1 pandemic from 2009/2010, it is worth highlighting our results for this particular season (Table 1, Fig. 2). The median values of the minimal and maximal basic reproduction number were 1.17 (interquartile range (IQR): 1.11–1.24) and 1.65 (IQR: 1.52–1.78), respectively, and were not significantly different to values from other seasons between 2003 and 2015 (Fig. 3). The most pronounced differences compared to the other seasons was the drop in the median values of susceptibility from over 88% for 0–4, 5–14, 15–29 and 30–64 year olds to 0.30 (IQR: 0.20–0.41) for 65+ year olds. Susceptibility for the oldest age group increased considerably in the subsequent season 2010/2011, which was expected because only 55% of influenza cases were attributed to H1N1 and 43% were attributed to Type B at that time (Hôpitaux Universitaires Genève, accessed 24 Nov, 2017). Lastly, ascertainability also clearly differed between age groups during the 2009/2010 season and was significantly lower for 30–64 and 65+ year olds compared to the three youngest age groups.

## 4. Discussion

We developed an influenza transmission model including age-structure and seasonal forcing. We then fitted the model to Swiss surveillance and virological test data from 2003-2015 using a Bayesian framework. The model was able to reproduce the transmission dynamics of ten influenza seasons and allowed us to infer critical parameters describing the transmission of and susceptibility to the annual epidemics. The median of the posterior distribution of the maximal basic reproduction number, *R*_0,max_, ranged from 1.46 to 1.81, which is in good agreement with the range of values that have been estimated for seasonal influenza epidemics in various countries (Biggerstaff et al., 2014). The median estimates of the susceptibility to influenza ranged from 29% to 98% for different age groups, and typically decreased with age. There was a slight negative correlation between susceptibility and ascertainability which also declined with age. Finally, our Bayesian modeling framework identified a considerable reduction in susceptibility to the H1N1 pandemic from 2009/2010 in the oldest age group. This finding is in agreement with the observation that adults over the age of 60 years had a higher frequency of cross-reactive antibodies against the pandemic strain, possibly due to prior exposure to antigenetically similar viruses (Hancock et al., 2009).

This is the first mathematical modeling study that investigates the transmission dynamics of influenza in Switzerland for several seasons. Our state-of-the-art Bayesian modeling framework and MCMC methods allowed us to indirectly infer critical parameters that describe the transmission of and susceptibility to influenza. Furthermore, our study also shows how virological test data can be used in combination with ILI data for producing more realistic models.

There are a number of limitations with our study. First, our model contains a relatively large number of free and fixed parameters that describe the influenza transmission dynamics in different age groups. Characteristically for our Bayesian modeling approach, the posterior distributions of our parameters estimates depend on the assumed prior distributions that were informed by the literature. Some of these parameters are relatively well-known and characterized, such as the infectious period or the range of *R*_0_. For others, we had to make some reasonable assumptions. Particularly for the ascertainability, which describes the probability that an influenza-infected individual becomes symptomatic, seeks a GP and would exhibit a positive virological test, little information is available. Due to the observed correlation between ascertainability and susceptibility in our model, it would be desirable to have a better informed prior distribution of the former parameter for determining the latter more precisely. The probability of ascertainability also affects the estimated influenza attack rates that vary between 2–56% for the different age groups (see Supplementary material). There exists considerable uncertainty about the annual attack rates for seasonal influenza (Somes et al., 2018), and estimates based on seroprevalence surveys are highly sensitive to seropositivity thresholds (Wu et al., 2014). While some of our values are relatively high, they are consistent with estimates of 2–4 influenza infections per decade at risk (Kucharski et al., 2015). Nevertheless, the relatively high estimates of the influenza attack rates for some age groups could indicate a higher ascertainability in Switzerland, compared to the study by Baguelin et al. (2013) for England and Wales, which would result in lower attack rates. Second, we did not consider detailed human demography of Switzerland (e.g., death of individuals) and relied on a simulated social contact matrix. We also ran our model using the German POLYMOD data from Mossong et al. (2008) (see Supplementary material) and found that this results in only minor differences in the model fits and posterior distributions. Hence, we believe that the reconstructed social contact matrices from Fumanelli et al. (2012), and similar approaches (Prem et al., 2017), provide a useful template for incorporating social contact data into mathematical models of influenza transmission when no survey data are available. Third, our model did not take into consideration the different virus strains/subtypes that were circulating during each season (Smieszek et al., 2011). Hence, we assumed that infection by strain/subtype provides immunity against infection by another strain/subtype. Furthermore, the epidemics of different strains/subtypes can peak at different times, and their transmission rates can differ between age groups. For example, during the 2012/2013 season, the 0–4 and 5–14 year olds were mainly infected with influenza type B, while 65+ year olds were primarily infected with H3N2 (Hôpitaux Universitaires Genève, accessed 24 Nov, 2017). Pooling the different strains/subtypes together is less problematic as long as the seasonal influenza epidemics are dominated by one strain/subtype. However, this is not the case for all seasons from 2003–2014 (see Supplementary material). Since the weekly numbers of positive virological samples in our data were often low, stratifying the model by strains/subtypes would have considerably limited the ability of our Bayesian modeling framework to infer the necessary parameters. This is why we decided to focus on the overall dynamics of influenza transmission and fitted the model for each season individually.

Finally, we did not have data on the yearly influenza vaccination uptake in the Swiss population. Hence, we could not investigate the effects of current or alternative vaccination strategies on influenza transmission in Switzerland. It is also important to note that the reduced levels of susceptibility among different age groups should be interpreted as a result of (cross-)immunity from previous influenza infections as well as vaccination.

Our modeling framework was inspired by the study from Baguelin et al. (2013) that analyzed strain-specific transmission of seasonal influenza in England and Wales from 1995/1996 to 2008/2009 including data from weekly virological swabs. The authors concluded that the efficiency of the traditional vaccination strategy targeting older adults and risk groups could be improved by including children. The relatively high incidence and susceptibility among young age groups (0–4 and 5–14 year olds) in our model suggests that it would yield similar results if children were vaccinated. In contrast to Baguelin et al. (2013), we included seasonal forcing in our model which improved the model fits of the peak incidence (Shaman et al., 2010, 2011; Lofgren et al., 2007) while not considerably increasing model complexity.

Lunelli et al. (2013) used a similar SEIR to better understand the influenza transmission dynamics in Italy during a 9-year period. Instead of virological data, they used annual serological data from antibody tests. This allowed them to estimate the susceptibility to influenza in each age group at the beginning and at the end of each season. While antibody tests have the advantage that they can be performed independently from a surveillance network, it is unclear how accurately they can be used as a marker for influenza infections in a population. The authors found reporting rates between 19.7% and 33.4%, which are considerably higher than our estimates of ascertainability as well as those estimated in the study by Baguelin et al. (2013).

Our modeling approach could be extended in several ways. Since we assumed seasonal forcing of influenza transmission, our model could in principle be fit to multiple seasons simultaneously (Goeyvaerts et al., 2015; Axelsen et al., 2014). Such models then allow to describe waning and boosting of immunity and can be used to predict the magnitude of outbreaks in upcoming seasons. Another possible extension would be to include more detailed contact structures, such as households (Tsang et al., 2016; Cauchemez et al., 2004) or social networks, that can affect transmission of influenza as well as the effect of control measures such as vaccination (Barclay et al., 2014).

This study shows how influenza surveillance and virological test data from Switzerland can be integrated into a Bayesian modeling framework. By assessing the underlying transmission dynamics of influenza, rather than just the incidence of ILI, the model complements current surveillance efforts and improves our understanding of seasonal influenza epidemics. Additional data, such as longitudinal antibody tests and surveys that study Swiss-specific social contact and mixing patterns as well health seeking behavior would help to further improve our model. While the presented modeling framework can be used to estimate the age-specific transmission of and susceptibility to past influenza epidemics, it would be desirable to incorporate vaccination data in future studies for assessing the effectiveness of current and alternative vaccination scenarios in Switzerland.

## Acknowledgments

We would like to thank Sentinella, the Swiss Sentinel Surveillance Network, and the Swiss Federal Office of Public Health (SFOPH) for providing the surveillance data, and the Virology Laboratory from the Geneva University Hospitals for providing the virological data.

